# Hierarchy of connectivity-function relationship of the human cortex revealed through predicting activity across functional domains

**DOI:** 10.1101/591776

**Authors:** Dongya Wu, Lingzhong Fan, Ming Song, Haiyan Wang, Congying Chu, Shan Yu, Tianzi Jiang

**Author notes:** Dongya Wu and Lingzhong Fan contributed equally to this work. **Corresponding Author:** Tianzi Jiang, Institute of Automation, Chinese Academy of Sciences, Beijing 100190, China.

## Abstract

Many studies showed that anatomical connectivity supports both anatomical and functional hierarchies that span across the primary and association cortices in the cerebral cortex. However, it remains unclear whether a hierarchy of connectivity-function relationship (CFR) exists across the human cortex as well as how to characterize the hierarchy of this CFR if it exists. We first addressed whether anatomical connectivity could be used to predict functional activations across different functional domains using multilinear regression models. Then we characterized the CFR by predicting activity from anatomical connectivity throughout the cortex. We found that there is a hierarchy of CFR across the human cortex. Moreover, this CFR hierarchy was correlated to the functional and anatomical hierarchy reflected in functional flexibility, functional variability, and the myelin map. Our results suggest a shared hierarchical mechanism in the cortex, a finding which provides important insights into the anatomical and functional organization of the human brain.

## Introduction

Converging evidence indicates that spatial–temporal hierarchical organizations that span between the primary and association cortices exist in the primate cerebral cortex^1, 2^. This hierarchy strongly affects both the functional and anatomical organization of the cortex. A spatial hierarchy has been found using the laminar patterns of anatomical projections^3^ and was further indicated by the macroscale cortical myelin estimated using magnetic resonance imaging (MRI)^4^. Recently, two parallel modeling studies also indicated that heterogeneity of the anatomical cortical organization shapes the large-scale hierarchies of neural dynamics^5, 6^. As a temporal counterpart, a hierarchy of different timescales across the cortex has also been observed and modeled based on invasive tract-tracing connectivity in the macaque neocortex^2^, which indicated the presence of a functional spectrum from early sensory to cognitive cortical areas. In other words, the various functions of the brain, from simple sensory-motor processing to complex cognitive processing, appear to be mediated in parallel with these anatomical and functional hierarchies. At the same time, the above studies indicate that extrinsic anatomical connectivity plays an important role in supporting such inter-areal hierarchical heterogeneity. In addition, it is well known that the function of a cortical region is constrained by the underlying extrinsic anatomical connectivity^7^. However, the extent that function is shaped by anatomical connectivity, which can be termed as the connectivity-function relationship (CFR), has not been fully characterized across the whole cortex. Moreover, whether there is a hierarchy in the CFR across different cortical areas remains unclear.

Anatomical connectivity is regarded as the basis for brain functions in the cortex^7^ and can provide a more powerful framework than physical location for describing brain functions^8^. Several studies on the characterization of CFR have already been made. The early studies investigated whether the boundaries of distinct brain regions characterized by anatomical connectivity coincide with boundaries of functionally distinct regions^9–12^. However, rich functional differentiations exist even within a region^13, 14^. Other researchers further suggested that variations in the connectivity profile can explain functional activations at a fine scale^7, 15^. Recent studies assessed the CFR by investigating the extent to which the functional activation of a few visual contrasts could be predicted from anatomical connectivity at a voxel-wise scale^16, 17^, but most of the brain regions that are activated in a few visual contrasts are only specialized for face processing or visual functions^18–22^.

However, it is well known that the primary sensory and motor cortices differ from the association cortices in terms of their laminar organization and afferent and efferent connections^23^. The primary cortices are organized in a topographic fashion, forming preferentially local networks, whereas the association cortices are organized in a widely distributed manner that yields complex zones^24, 25^. The existence of anatomical and functional organizational properties that differ between the primary and association cortices may indicate that the CFR in the association cortices differs from that in the primary cortices. Therefore, whether anatomical connectivity can predict functional activations to the same degree along the cortical hierarchy is still unknown, especially in the association cortices, which are functionally flexible^26–28^ and highly variable across individuals^29^. Furthermore, it remains unclear how this inter-areal heterogeneity in the connectivity organization affects the CFR.

The current study addressed the above questions by investigating the relationship between anatomical connectivity and functional activation in different regions along the cortical hierarchy, based on the Brainnetome atlas^30^. We used a prediction model to assess the CFR to ensure that the CFR captured by the multilinear model was not a result of over fitting. We adopted the Human Connectome Project (HCP) dataset, which includes seven functional domains, for two reasons: First, we aimed to test whether the CFR would be consistent across different task states. Second, because different tasks might activate different cortical regions, we wanted to include as many cortical regions as possible. Finally, to determine whether the pattern of the CFR had a specific meaning, we investigated whether the pattern of the CFR was also reflected in the functional and anatomical hierarchy of the cortex.

## Results

### Comparison between actual activation and predicted activation

Since the CFR was assessed via the similarity between predicted activation and actual activation, we first investigated the anatomical connectivity’s predicted activation visually. We selected several representative contrasts and plotted the actual and predicted activation of subject 120111 on the brain surface in Fig. 1; more examples are shown in Fig. S1. The threshold values for the individual activation maps were based on a Gaussian-two-gamma mixture model^31, 32^, where the Gaussian represented the distribution of noise and the two gammas represented the distributions of the positive and negative activations. The positive and negative thresholds were chosen to be the medians of the two gammas. The overall pattern of the predicted activation was very similar to that of the actual activation. The similarities (evaluated by correlation) are quantitatively presented in the third column of Table S2. Anatomical connectivity predicted the overall patterns of task activation in various functional domains. To ensure that the prediction results were not completely driven by the parcellation adopted, a direct quantitative comparison of the predictions based on the Brainnetome atlas^30^ and the HCP_MMP1.0 parcellation^33^ is shown in the fourth column of Table S2. The similarities (evaluated by correlation) between the predicted activations based on the Brainnetome atlas and on the HCP_MMP1.0 parcellation were very high; thus the parcellation scheme had little influence on the prediction results.

**Figure 1.**
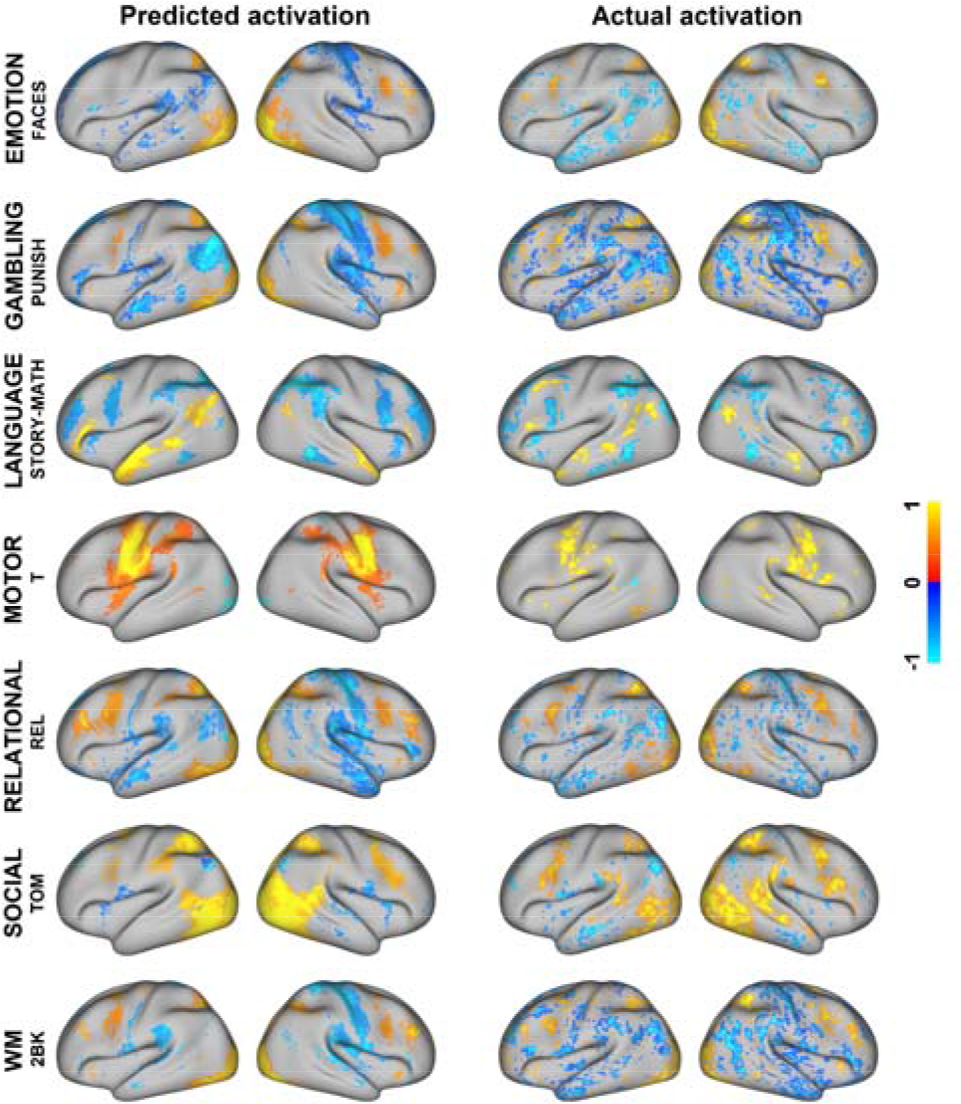
Comparison between the actual and predicted activations. The left column represents the predicted activation and the right column represents the actual activation. Different rows represent activations of different contrasts. We mapped the 98 percentiles of the most positive and negative data values to 1 and −1. The overall patterns of the predicted activation were very similar to those of the actual activation.

### Statistical tests of the CFR

The previous section showed that anatomical connectivity predicted the overall pattern of task activation, but to compare the CFR for each region, we must first examine the CFR of each region statistically. Statistical analyses using a permutation test that shuffled the parings between connectivity features and task activations were performed to assess the reliability of the connectivity model. If the mean prediction accuracy of the connectivity model was higher than the 95th percentile of the mean prediction accuracy of the random models, we regarded the connectivity model as statistically better than random. We calculated the number of regions that had connectivity models statistically better than random (Table S3). We found that the CFR was statistically better than random in many regions across most contrasts. We did two sample *t*-tests and found that in all the contrasts the regions that had predictions better than random had task activations (defined in the methods part) significantly higher (*p* < 1e-3) than regions that had predictions that were not better than random. The CFR for a few contrasts, such as PUNISH-REWARD, were better than random in only a few regions because these contrasts had a very low activation level throughout the whole cortex.

Additionally, distance may influence the connectivity model since regions that are closer together are more likely to be connected and co-activated. Therefore, an additional control analysis using a distance model was conducted to ensure that the performance of the connectivity model was not driven by the spatial relationships. Instead of using the connectivity strength of various vertices to other brain regions, the distance model used the Euclidian distance from the vertices to the center of other brain regions as features. A comparison of the prediction accuracy between the distance model and the connectivity model is shown in Fig. S2. Anatomical connectivity had a better prediction accuracy than the distance model. After regressing out distance from the connectivity profile, anatomical connectivity still had a prediction accuracy that was better than random in many regions (Table S3).

### Hierarchy in the CFR

The previous section indicated that the CFR was statistically meaningful across different domains, but regional differences in the CFR still existed. We investigated the regional differences based on the Brainnetome atlas and on the seven functional networks^25^ (See Fig. 2). We mapped each of the 210 cortical regions to one of the seven functional networks that achieved maximum overlap, and the CFRs in each of the seven functional networks were averaged. As shown in the previous section, the CFR was not statistically significant in regions that had a low task activation; thus we only included regions that had a task activation that was higher than a given threshold. The calculation of the task activation and the determination of the threshold are provided in the Methods section. The results show that the CFR in the sensory-motor networks was stronger than that in the association networks, with an exception that the CFR in the visual network was weak under the language story contrast because this task only included an auditory stimulus but not a visual stimulus. Since different task contrasts activated different brain regions, we could not directly compare the CFR between two regions that had totally different activation levels in the same contrast, such as between the motor and prefrontal brain regions in the motor contrast. Therefore, to allow for the comparison of the CFR between any two regions, we focused on the consistency of the CFR in all the contrasts instead of in a single one and averaged the CFR across all the contrasts (in Fig. 3), including only the activated regions (the same regions as in Fig. 2) in the averaging process. To assure that this selective averaging result was not dramatically driven by the threshold level, we verified that the result was consistent across different threshold levels (Fig. S3). We found that a hierarchy existed in the CFR: the CFR was relatively high in the sensory-motor networks, moderate in the dorsal and ventral attentional networks, and relatively low in the frontoparietal, default, and limbic networks. The sensory-motor networks included regions such as the motor, auditory, and visual cortices, and the association frontoparietal and default networks included regions such as the lateral prefrontal cortex and temporal-parietal junction. The CFR in the sensory-motor networks was significantly (*p*<1e-6) higher than that in the association networks under a two sample *t*-test.

**Figure 2.**
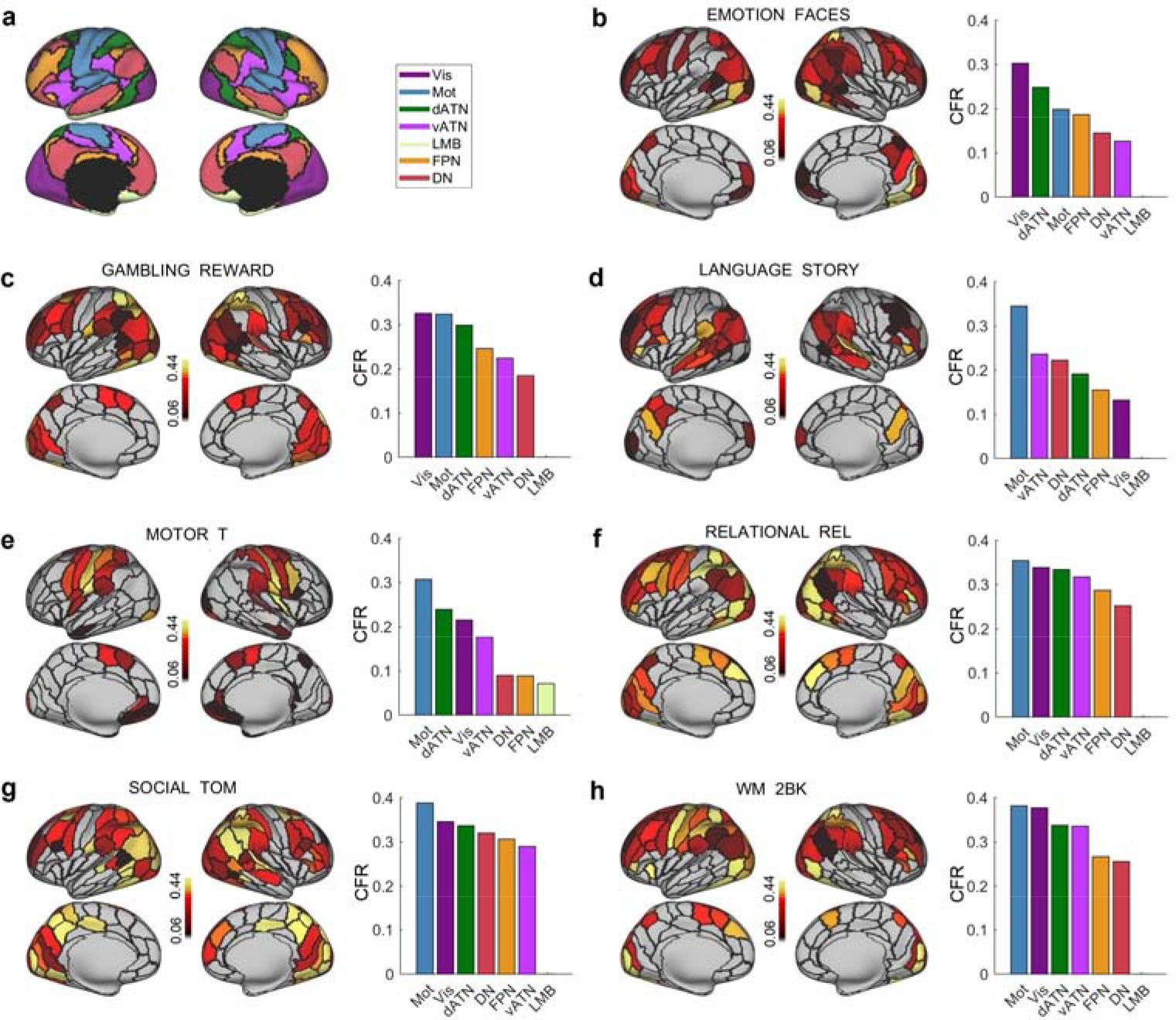
Spatial distribution of the CFR in different contrasts. (a) The seven functional networks of the cortex. The colors represent the mapping of the seven networks, namely the frontoparietal control (FPN), ventral and dorsal attention (vATN, dATN), default (DN), and limbic (LMB) networks that constitute the association networks and the motor-auditory (Mot) and visual (Vis) networks that constitute the sensory-motor networks. The spatial distribution of CFR and the rank of the CFR in the seven networks are shown in different subplots, with (b) EMOTION FACES, (c) GAMBLING PUNISH, (d) LANGUAGE STORY, (e) MOTOR T(Tongue), (f) RELATIONAL REL, (g) SOCIAL TOM, and (h) WM 2BK.

**Figure 3.**
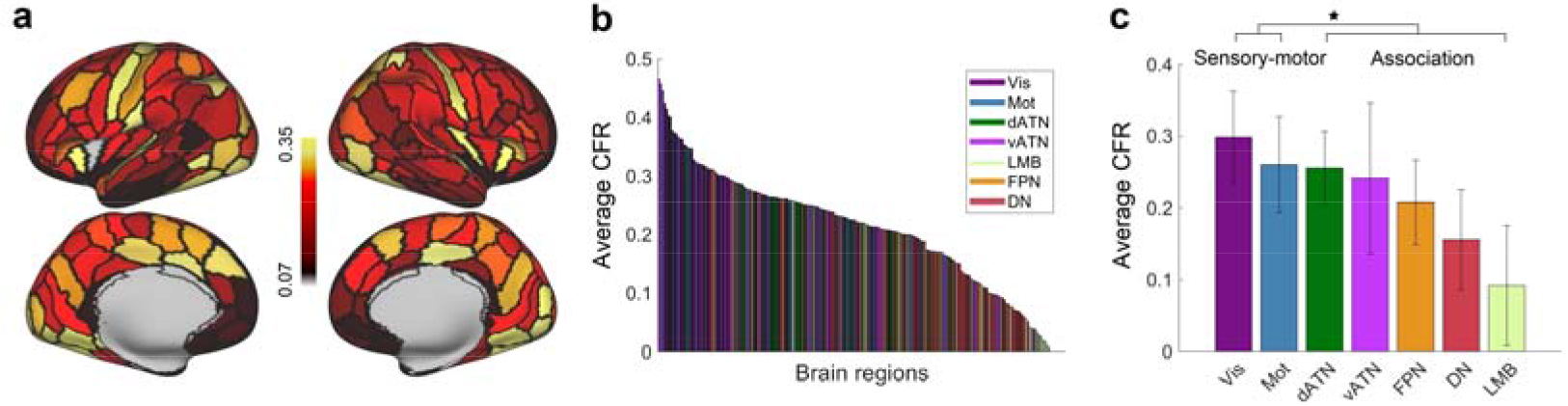
Spatial distribution of the average CFR. (a) Spatial distribution of an average CFR in the Brainnetome atlas. (b) The average CFR was ranked, and the color represented the mapping of the seven networks. The CFR was highest in regions that are located mostly in the Vis and Mot networks. (c) Rank of the average CFR in the seven networks. The star symbol indicates that the CFR in the sensory-motor networks was significantly (*p*<1e-6) higher than that in the association networks under a two sample *t*-test.

### Hierarchy in the CFR was related to functional flexibility

We revealed the hierarchy in the CFR in the previous section. Next, we explored the connectivity features that contributed to the CFR in the multilinear regression model. We extracted the significant (*p*<1e-4) connectivity features of each cortical region in predicting each task’s activations, and these connectivity features were further used to investigate the functional flexibility of each region. The functional flexibility was defined as the variation in connectivity features across different tasks, which was to assess whether a cortical region utilized similar or different patterns of connectivity features in predicting its task activations across different tasks. The calculation of functional flexibility is provided in detail in the Method section, and the same threshold levels as in the previous section were used to identify the activated regions in each task. Examples of two regions are presented in Fig. 4. The region A37lv (in the fusiform, lateroventral Area 37) had a flexibility of 0.37 and had relatively similar patterns of connectivity features across different tasks, and the region A46 (in the lateral prefrontal cortex, Area 46) had a flexibility of 0.63 and had relatively different patterns of connectivity features across different tasks. The functional flexibility was also assessed in the seven functional networks, and the flexibility of the sensory-motor networks was significantly (*p*<1e-6) lower than that of the association networks under a two sample *t*-test. Further, the flexibility of each region was negatively correlated (*p*<1e-6) with the average CFR. Therefore, functional flexibility exhibited a hierarchy that was the inverse of the hierarchy in the CFR.

**Figure 4.**
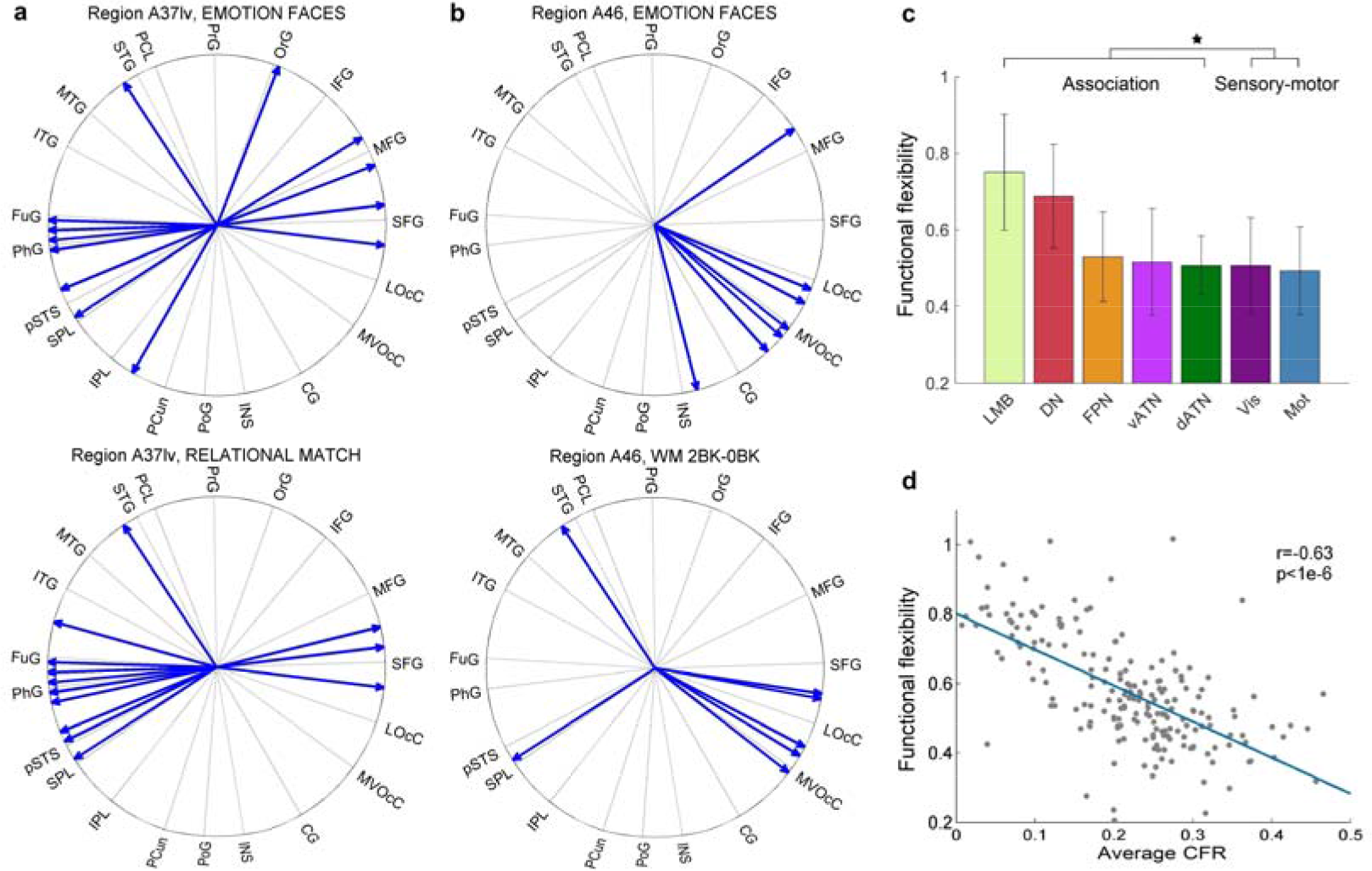
The flexibility of the connectivity features was related to the CFR. (a) The blue arrows represent the significant connectivity features of region A37lv (in the fusiform, lateroventral Area 37) in two tasks. The cortical region name is viewed in an anti-clockwise manner, with each name labeling the following cortical regions in the block. The abbreviations are included in the Methods section. (b) The blue arrows represent the significant connectivity features of region A46 (in the lateral prefrontal cortex, Area 46) in two tasks. (c) The functional flexibility was ranked in the seven functional networks. The star symbol indicates that the flexibility of the sensory-motor networks was significantly (*p*<1e-6) lower than that of the association networks under a two sample *t*-test. (d) The flexibility of each region was negatively correlated (*p*<1e-6) with the average CFR.

### Hierarchy in the CFR was related to functional variability

After revealing that the hierarchy in the CFR was related to functional flexibility, we investigated whether the hierarchy in the CFR was related to another index of functional hierarchy of the cortex. We explored each region’s inter-subject task variation, as a previous study indicated that the inter-subject functional variability reflected a hierarchical architecture of the cortex^29^, and we used task activation as a control variable to ensure that the result was not a byproduct of the activation level. Each region’s task activation profile and inter-subject task variation are plotted in Fig. 5, with the same threshold method used as that in Fig. 2; all regions are color coded by their CFR. The CFR was relatively lower in regions with a larger inter-subject task variation and a lower task activation, and the CFR was relatively higher in regions with a smaller inter-subject task variation and a higher task activation. We quantitatively calculated the amount of variance in the CFR that could be explained by the task activation or the inter-subject task variation, and the results are provided in Table 1 and Table S4. Task activation was positively correlated with the CFR across all contrasts, whereas inter-subject task variation was negatively correlated with the CFR. However, task activation could barely explain any of the variance in the CFR after the inter-subject task variation was regressed out. These results indicated that the CFR was mainly dominated by inter-subject task variation: the CFR was high in regions with a small task variation and was low in regions with a large task variation. The spatial distribution of the average inter-subject task variation across all contrasts (calculated in the same way as the average CFR) is provided in Fig. 6. The inter-subject task variation had an inverse hierarchy to the CFR and was negatively correlated with the CFR; the inter-subject task variation was lowest in the sensory-motor networks, modest in the dorsal and ventral attentional networks, and highest in the frontoparietal and default networks.

**Table 1.** Quantitative analysis of the factors that affected the CFR. CFR-variation represents the CFR after the inter-subject task variation was regressed out, and similarly for the CFR-activation. The plus sign indicates positive correlation with the CFR, and the minus sign indicates negative correlation. Task activation was positively correlated with the CFR, whereas inter-subject task variation was negatively correlated with the CFR. Task variation explained significantly (*p* < 1e-6) more variance in the CFR than task activation did under a paired-t test across all contrasts. Task activation could barely explain any of the variance in the CFR after the inter-subject task variation was regressed out. The results for all contrasts are shown in Table S4.

| Contrasts | Task activation(+) |  | Inter-subject task variation(-) |  |
| --- | --- | --- | --- | --- |
|  | CFR | CFR-variation | CFR | CFR-activation |
| EMOTION FACES | 0.16 | 0.05 | 0.60 | 0.29 |
| GAMBLING REWARD | 0.13 | 0.04 | 0.59 | 0.33 |
| LANGUAGE STORY | 0.19 | 0.01 | 0.67 | 0.38 |
| MOTOR T | 0.32 | 0.05 | 0.67 | 0.17 |
| RELATIONAL REL | 0.04 | 0.07 | 0.58 | 0.46 |
| SOCIAL TOM | 0.05 | 0.03 | 0.53 | 0.40 |
| WM 2BK | 0.07 | 0.06 | 0.56 | 0.39 |

**Figure 5.**
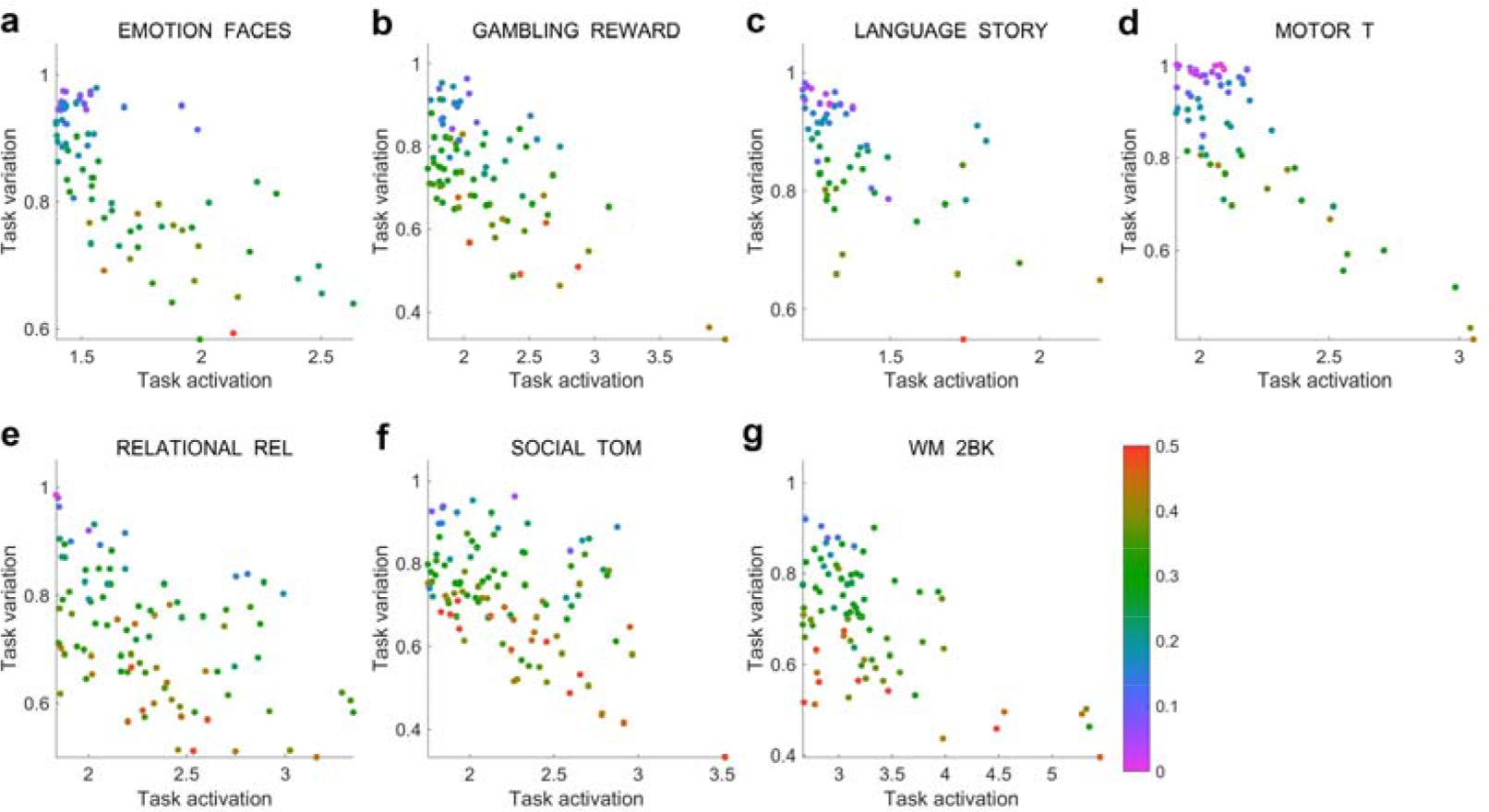
Analysis of the factors that affected the CFR. Each region’s task activation profile was plotted against that region’s inter-subject task variation, and all the regions are color coded according to their CFR. The CFR was low in the regions that had a large inter-subject task variation and a low task activation and was high in the regions that had a small inter-subject task variation and a high task activation. Different subplots represent the results of different contrasts, with (a) EMOTION FACES, (b) GAMBLING REWAED, (c) LANGUAGE STORY, (d) MOTOR T, (e) RELATIONAL REL, (f) SOCIAL TOM, and (g) WM 2BK.

**Figure 6.**
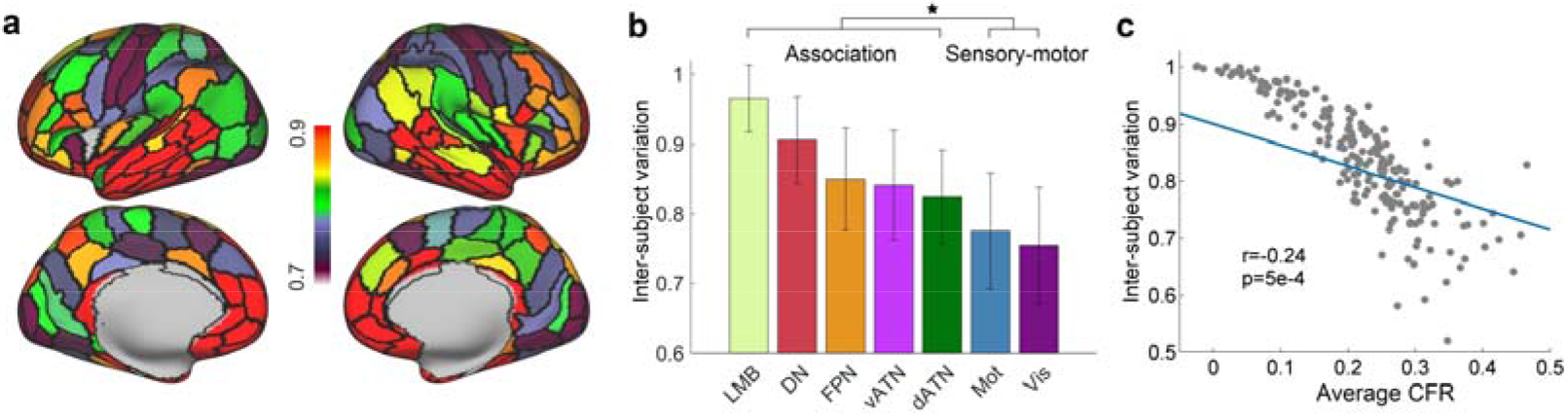
Spatial distribution of the inter-subject task variation. (a) Spatial distribution of the average inter-subject task variation in the Brainnetome atlas. Regions such as the lateral prefrontal cortex and temporal-parietal junction had large inter-subject variations, and regions such as the motor and visual cortices had small inter-subject variations. (b) The rank of the inter-subject task variation in the seven networks. The sensory-motor networks had small inter-subject variations, and the association networks had large inter-subject variations. The star symbol indicates that the inter-subject variation of the sensory-motor networks was significantly (*p*<1e-6) lower than that of the association networks under a two sample *t*-test. (c) The inter-subject task variation was negatively correlated with the CFR.

### Hierarchy in the CFR was related to the myelin map

To test whether the hierarchy in the CFR was related to the anatomical hierarchy of the cortex, we compared the pattern of CFR with that of myelin, as a previous study indicated that the myelin map reflects the anatomical hierarchy of the cortex^4^. The spatial distribution of the myelin map is shown in Fig. 7. The myelin map was an average result from all subjects in the HCP S1200 release. A similar pattern of hierarchy in the CFR was also exhibited in the myelin map. The myelin map exhibited significantly (*p*<1e-6) higher values in the sensory-motor networks than in the association networks, and the myelin map was significantly (*p*<1e-6) correlated with the CFR. These results indicate that the hierarchy in the CFR is also a reflection of the anatomical hierarchy of the cortex.

**Figure 7.**
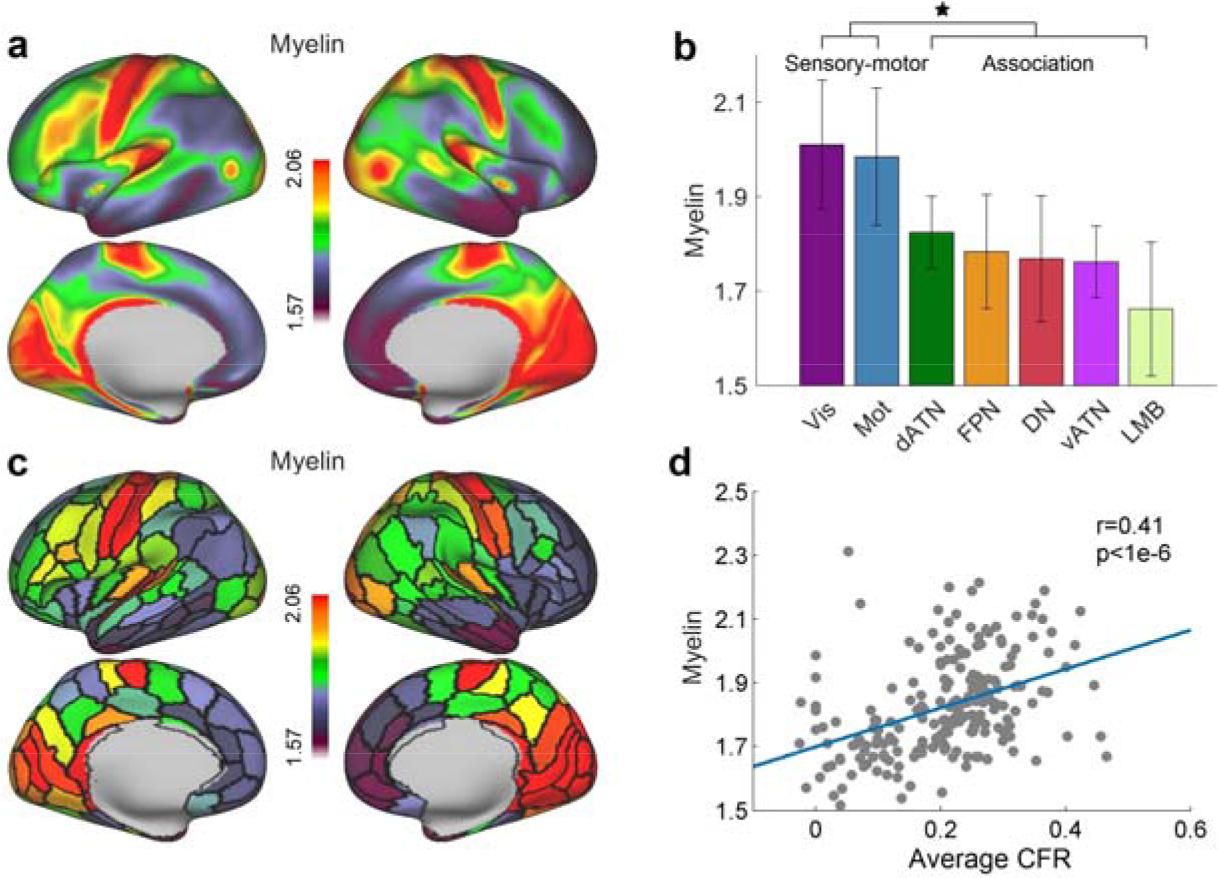
Spatial distribution of the myelin map. (a) Spatial distribution of an average myelin map on the cortex. (b) Rank of the seven functional networks in the myelin map. The star symbol indicates that the myelin in the sensory-motor networks was significantly (*p*<1e-6) higher than that in the association networks under a two sample *t*-test. (c) Spatial distribution of an average myelin map on the Brainnetome atlas. (d) The myelin was positively correlated with the average CFR based on the Brainnetome atlas.

## Discussion

In the present study, we investigated whether a cortical hierarchy of the CFR exists throughout the cortex. Specifically, our results showed that the CFR was statistically better than random models in most regions across seven functional domains. Moreover, we revealed that the CFR in the sensory-motor networks was higher than the CFR in the association networks and that similar regional differences between the sensory-motor and association cortices were also reflected in the organization of functional flexibility, functional variability, and the myelin map, a finding which suggests that the CFR has a hierarchical structure throughout the cortex.

The CFR was assessed via the predictive ability of anatomical connectivity, and the rationality of this assessment was confirmed by the following points. Since the connectivity model outperformed the random control model in most regions across seven functional domains, the statistical significance of the CFR was confirmed in most cases. The regions in which the CFR failed to pass the statistical test had statistically lower activation levels than the regions in which the CFR passed the statistical test. Therefore, testing the CFR in task-relevant regions is necessary, so we adopted an HCP task fMRI dataset that incorporates a wide range of functional domains to cover as many region’s functions as possible. Previous studies have already showed that a close relationship exists between anatomical connectivity and visual functions via the predictive ability of anatomical connectivity^16, 17^. Our result demonstrates that this CFR is not restricted to visual functions and that brain function is closely related to anatomical connectivity from the primary sensory and motor functional domain to the high order cognitive functional domain, a finding which supports the assessment of CFR in high order functions.

Moreover, we also revealed that the CFR was hierarchical across the whole cortex. The CFR was relatively high in the sensory-motor and visual networks, moderate in the attentional networks, and relatively low in the frontoparietal and default networks. This pattern of hierarchy in the CFR was further found to be negatively correlated with the functional flexibility of the anatomical connectivity profile. While the anatomical connectivity substrate is fixed, the functional repertoire of the brain is diverse. This kind of many-to-one mapping from function to structure suggests that a fixed structure can support flexible functions, and directly leads to cognitive degeneracy^34, 35^. Our result indicated that the association cortex is more flexibly involved in a variety of tasks than is the sensory-motor cortex, coinciding with a review that suggested that the divergence between structure and function is enabled more at the global integration level^36^. This many-to-one mapping between function and anatomical connectivity may make the prediction of function from anatomical connectivity more difficult in the association cortex. In addition, hierarchy in the CFR was found to be negatively correlated with inter-subject functional variability and to be positively correlated with the myelin map. Because functional flexibility and functional variability were higher in the association cortex than in the sensory-motor cortex and the myelin map was negatively correlated with the anatomical hierarchical level of the cortex^4^, the above finding suggests that the CFR decreases along both the functional and anatomical hierarchical axes from the sensory-motor to the association cortex. Because the attentional networks are spatially located between the sensory-motor-visual networks and the frontoparietal-default networks, the hierarchical structure in the CFR is very similar to the large-scale hierarchical gradients^37^ that span between the sensory-motor and association areas in various cortical organizations. For example, a previous study showed that concrete-to-abstract semantic gradients from the sensory-motor to the association cortices exist in the functional processing hierarchies^38^. In addition the gradients in functional processing hierarchies were likely to be supported by the gradients in connectivity^39^ and microstructure such as myelin^40^ and cortical thickness^41^, which were also likely to have a genetic basis in that gene expression has been shown to separate the sensory-motor cortices from the association cortices^42^. Therefore, our finding may suggest a hierarchical gradient in the CFR that has genetic and microstructural bases.

The consistency between the hierarchy in the CFR with both the functional and anatomical hierarchies of the cortical organization implies a shared mechanism in the hierarchical structure of the human brain. A plausible explanation as to why these evidences are tied together is that: compared with the sensory-motor cortex, the association cortex, which is late-developing in evolution^43^ and human development^44^ and thus possesses more variability^29^, is essential to more complex brain functions^24^ and acts as a hub of integration^45^, thus flexibly participating in multiple functions. The high functional load in the association cortex may require greater expansion^46, 47^ and more complex structural substrates. For example, the association cortex exhibits more dendritic branching complexity and dendritic spine density^48^, which also influences the neuronal functional properties^49^. Using the myelin map as a proxy, Burt et al. verified that the number of spines on pyramidal cell dendrites increases along a hierarchical axis from the sensory to the association cortex^4^, thus endowing the association cortex with more recurrent synaptic excitation in local cortical microcircuits supporting cognitive computations^50^. Therefore, the complex functions in the association cortex cannot be solely attributed to the source of their inputs determined by extrinsic connectivity^48^, and extrinsic anatomical connectivity alone may explain the relatively fewer functional activations in the association cortex due to the more complex structural substrates that underlie its functions, suggesting that more microstructural properties should be considered when studying the complex functions of the association cortex. This combination of factors behind the hierarchy in the CFR provides important insights into the understanding of the anatomical and functional organization of the human brain.

In conclusion, we verified that the brain functions were constrained by anatomical connectivity heterogeneously across the cortex and revealed that the hierarchical structure in the CFR was related to both the functional and anatomical hierarchies in cortical organizations. We provided an extensive delineation of the relationship between functional activation and anatomical connectivity in the whole cerebral cortex across various functional domains. Investigating the cortical function with respect to anatomical connectivity improves our understanding of how the cortical function emerges from connectivity constraints. Future work can build more complex models that incorporate microstructural properties to better characterize brain functions.

## Methods

### Human Connectome Project data

We used the minimally pre-processed data^51^ provided by the HCP. We randomly selected 100 unrelated subjects from the S1200 release. The information about the 100 unrelated subjects is listed in Table S1. Since we performed the analysis at the vertex level rather than at the subject level, the number of samples was far more than 100; these 100 subjects were sufficient to get a reliable prediction result. See Fig. S4 for the supporting evidence.

Acquisition parameters and processing were described in detail in several publications^52–55^. Briefly, diffusion data were acquired using single-shot 2D spin-echo multiband echo planar imaging on a Siemens 3 Tesla Skyra system^54^. These consisted of 3 shells (b-values = 1000, 2000, and 3000 s/mm^2^) with 270 diffusion directions isotropically distributed among the shells, and six b = 0 acquisitions within each shell, with a spatial resolution of 1.25 mm isotropic voxels. Each subject’s diffusion data had already been registered to his or her own native structural space^51^. Task fMRI scans were acquired at 2 mm isotropic resolution with a fast TR sampling rate at 0.72 s using multiband pulse sequences^55^.

We used the task fMRI data that were projected into 2 mm standard CIFTI grayordinates space, and the multimodal surface matching (MSM) algorithm^56^ based on areal features (MSMAll) was used for accurate inter-subject registration. The task fMRI contained 86 contrasts from seven task domains, labeled as EMOTION, GAMBLING, LANGUAGE, MOTOR, RELATIONAL, SOCIAL, and WM (working memory). The details of the tasks were described in Barch et al.^52^. Most of the contrast maps were paired with a related negative contrast, which amounted to adding a minus sign to the dependent variable in a multilinear model and was redundant for the purpose of regression modeling since it did not change the model performance. We excluded the redundant contrasts and kept 47 contrasts for further regression analysis.

### Anatomical connectivity profile

We calculated anatomical connectivity based on the Brainnetome atlas^30^, which contains 210 cortical regions in both hemispheres. Each of the 210 cortical regions was used as a seed region. For every seed region, therefore, there were 209 target cortical regions. All the vertices within the seed region were characterized by the anatomical connectivity of the 209 dimensions, representing the connectivity of each vertex in the seed region to the remaining 209 target regions. The anatomical connectivity was determined via probabilistic diffusion tractography. The white matter surface mesh aligned using MSMAll was used as the seed. Fiber orientations were estimated per voxel (three fibers per voxel), and probabilistic diffusion tractography was performed using FSL-FDT^57^ with 5000 streamline samples in each seed vertex to create a connectivity distribution to each of the remaining 209 target regions, while stopping tracking at the pial surface.

### Model training

To avoid overfitting, we separated the 100 subjects into a training group of 80 subjects and a testing group of 20 subjects. We only included the significant (*p*<1e-4) connectivity features on the training group for further prediction. The testing group was used to assess the training model. For each of the 47 contrasts, we performed a regression analysis on each of the 210 cortical regions using anatomical connectivity. The regression analysis was modeled as:Y = Xβ + E, where Y is the *z*-statistical value of the contrast maps and the mean of Y is subtracted to remove the intercept term; X represents anatomical connectivity; β is the regression coefficient to be estimated from the regression model. To train the regression model on the *i*-th cortical region, we concatenated all the vertices of the *i*-th cortical region into a column across the training subjects. Assuming that the *i*-th cortical region has *n*_*i*_ vertices, Y is a single column vector of length *N*_*i*_ = 80 × *n*_*i*_ representing the functional activation of all the seed vertices, and X is a matrix of *N*_*i*_ rows and 209 columns representing the connectivity features of all the seed vertices to the remaining 209 target regions, β is a single column vector of length 209, representing how each connectivity feature contributed to predicting a seed region’s functional activation. Similarly, we obtained a connectivity feature from the testing group. After estimating the regression model’s coefficients from the training group, we applied these coefficients to the testing group’s connectivity feature to get the predicted functional activation of the subjects in the testing group. To get the predicted functional activation for the entire cortex, we repeated the same procedure for every cortical region and then concatenated every region’s prediction.

### Assessment of the CFR

The CFR was assessed using the prediction accuracy of each testing subject’s functional activation to avoid overestimation on the training data. We correlated the predicted activation of every subject in the testing group with the actual functional activation of the same subject to evaluate the accuracy of the predictions, and the prediction accuracies of all the testing subjects were further averaged. We used the correlation coefficients (*r*) to assess the CFR instead of the mean squared error (MSE) or mean absolute error (MAE) because, unlike MSE and MAE, which are not standardized and un-bounded, *r* is standardized and bounded between 0 and 1. In addition, under the least square conditions of the regression model, the square of the correlation coefficient equals the proportion of the variance in the functional activation that can be explained by the connectivity features.

### Calculation of task activation and threshold

Each region’s task activation profile was calculated by first averaging each vertex’s absolute activation within a subject and then averaging each subject’s mean activation. We removed some regions that had a low-activation level in the process of averaging the CFR or inter-subject task variation within each of the seven functional networks. The threshold was determined by first estimating the density distribution of all regions’ task activations in each contrast using the “ksdensity” command in Matlab. Next, the density peak of the distribution was found, and finally the right endpoint of the 5% interval of the density peak was chosen to be the threshold for removing regions that had a relatively low activation level in each contrast.

### Calculation of inter-subject variation and functional flexibility

We calculated each region’s inter-subject task variation and the functional flexibility of each region in the same manner. We assessed the similarity of the task activation or connectivity feature between two subjects or two tasks by first reshaping the activation or connectivity feature into a column vector and then calculating the Pearson correlation. The variation in the activation or connectivity feature was calculated by averaging the similarities between all pairs of training subjects or tasks and then subtracting from one. If the variation in the task activation or connectivity feature was small, then the average similarity was close to one and the inter-subject variation or functional flexibility was close to zero, indicating a small inter-subject variation or flexibility; otherwise, if the average similarity was close to zero and the inter-subject variation or functional flexibility was close to one, the result indicated a large inter-subject variation or flexibility.

### Permutation test

We did 1000 random permutations to test the performance of the connectivity model statistically. We trained the models in the same manner, but the pairings between each vertex’s connectivity feature and its functional activation were shuffled. We then tested how these random models performed on the testing group. We got one mean prediction accuracy for the connectivity model and 1000 mean prediction accuracies for the random models. Then we calculated whether the mean prediction accuracy of the connectivity model was higher than the 95th percentile of the mean prediction accuracy of the random models to test whether the performance of the connectivity model was statistically meaningful. One thing to notice here was that we only shuffled the data in the training group but not in the testing group. Since we had trained the regression model one region at a time, we shuffled the pairings within the seed region but not across the whole cortex.

## Abbreviations of cortical labels

SFG: Superior Frontal Gyrus
MFG: Middle Frontal Gyrus
IFG: Inferior Frontal Gyrus
OrG: Orbital Gyrus
PrG: Precentral Gyrus
PCL: Paracentral Lobule
STG: Superior Temporal Gyrus
MTG: Middle Temporal Gyrus
ITG: Inferior Temporal Gyrus
FuG: Fusiform Gyrus
PhG: Parahippocampal Gyrus
pSTS: posterior Superior Temporal Sulcus
SPL: Superior Parietal Lobule
IPL: Inferior Parietal Lobule
Pcun: Precuneus
PoG: Postcentral Gyrus
INS: Insular Gyrus
CG: Cingulate Gyrus
MVOcC: MedioVentral Occipital Cortex
LOcC: Lateral Occipital Cortex

## Author contributions

Jiang, Fan, Yu, and Song designed and supervised the experiments. Wu, Wang, and Chu analyzed the data. All authors contributed to the writing of manuscript.

## Availability of code

The code for implementing all the analyses is available upon request.

## Acknowledgement

The authors thank Prof. Yuanye Ma and Prof. Simon B Eickhoff for their constructive suggestions which improved the manuscript and appreciate the editing assistance of Rhoda E. and Edmund F. Perozzi, PhDs.

