## supplementary material for "Hierarchy of connectivity-function relationship of the human cortex revealed through predicting activity across functional domains"

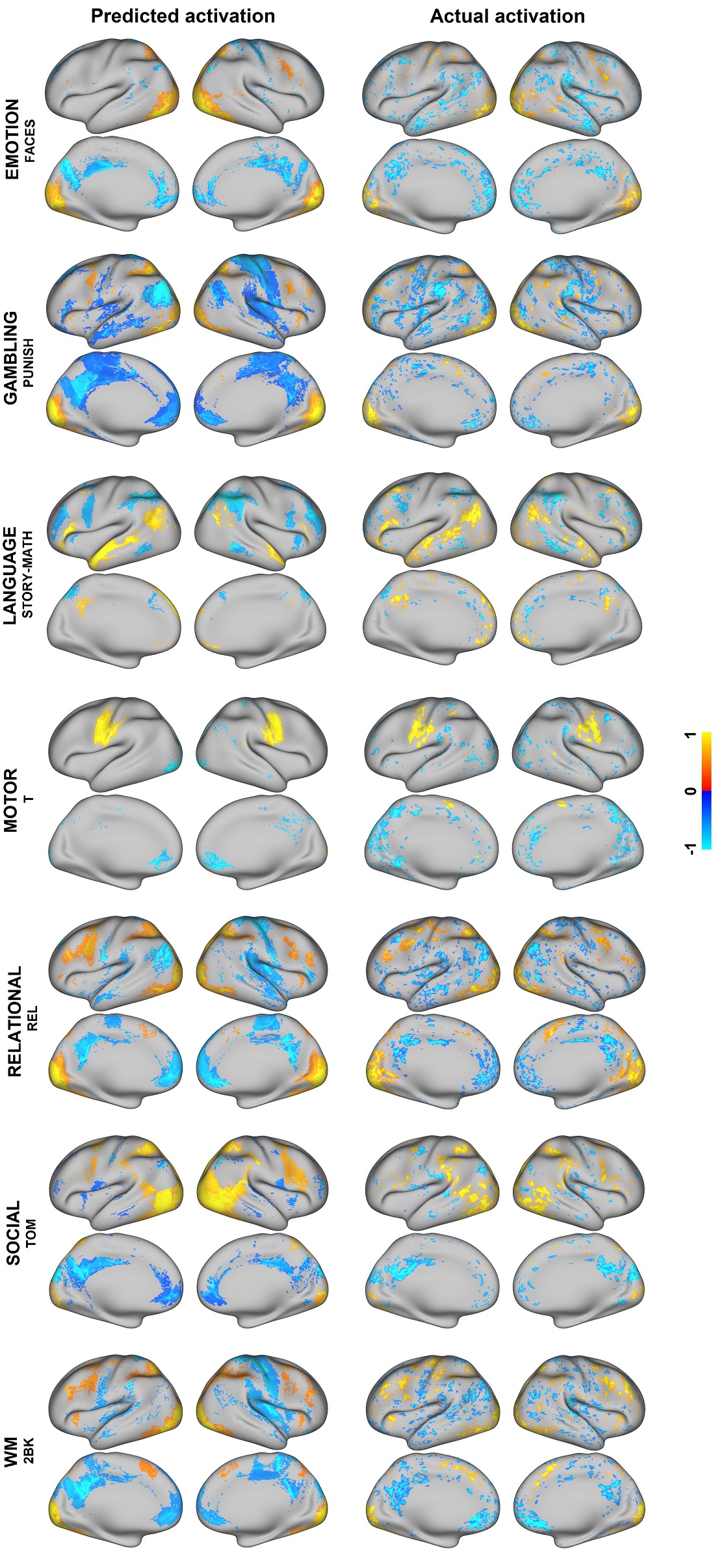

**Figure S1. Comparison between actual activation and** **predicted activation**. We selected several contrasts and formed a visualization of the actual and predicted activation of several testing subjects on the cortex. The left column represents the predicted activation made by anatomical connectivity and the right column represents the actual activation. We mapped the 98 percentiles of the most positive and negative data values to 1 and -1. The overall pattern of the predicted activation was very similar to that of the actual activation. Different rows represent activation of different subjects, with EMOTION FACES 113619, GAMBLING PUNISH 122317, LANGUAGE STORY-MATH 100408, MOTOR T 129028, RELATIONAL REL 899885, SOCIAL TOM 857263, WM 2BK 124422.

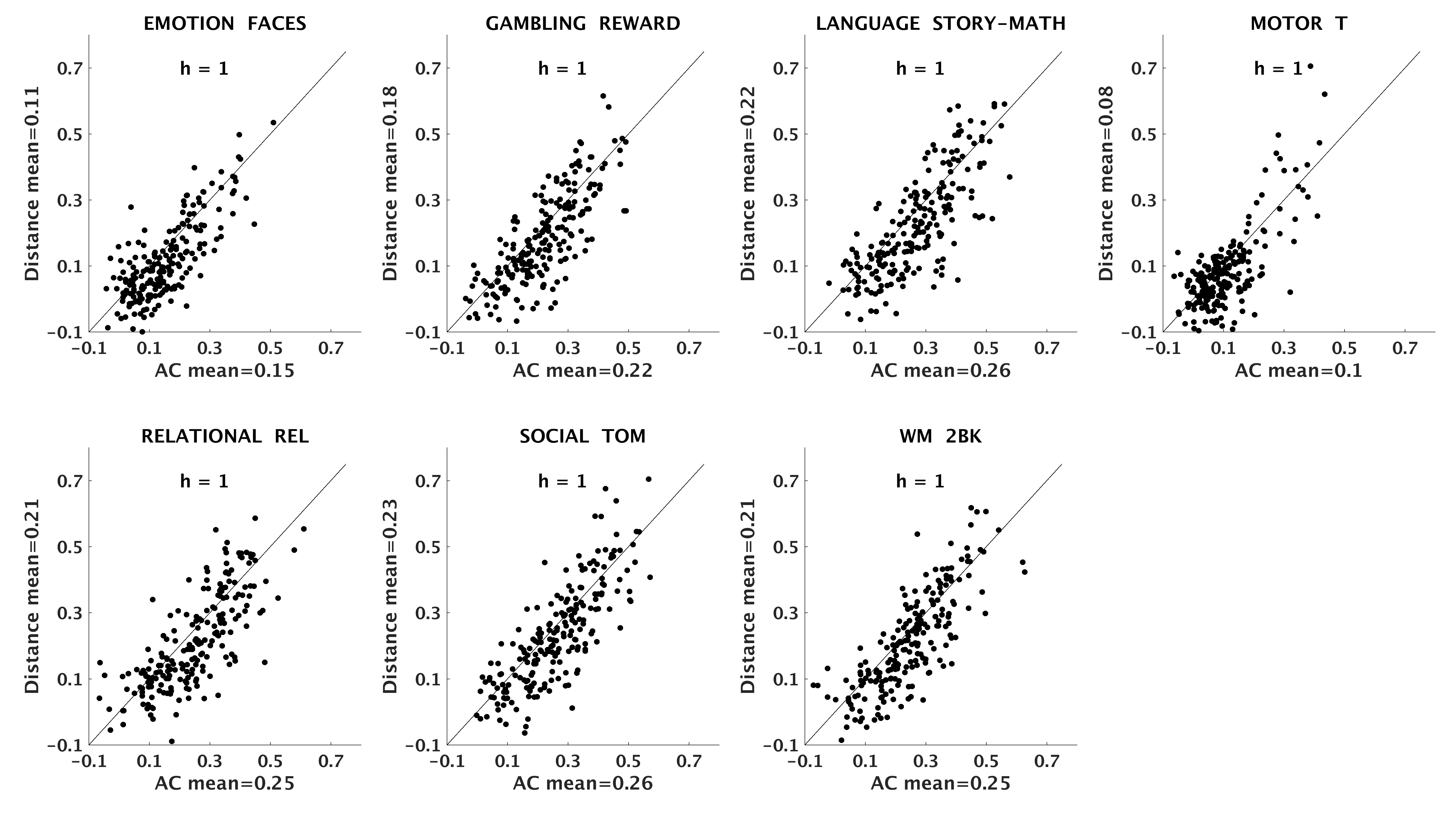

**Figure S2. Comparison of prediction accuracy between connectivity and distance model.** “h” indicated whether the difference in the prediction accuracy was significant (*p*<1e-4) under a paired-*t* test. The diagonal indicated that the prediction accuracy made by anatomical connectivity (AC) and that made by distance were equal. Anatomical connectivity had prediction accuracies statistically better than distance model in all contrasts.

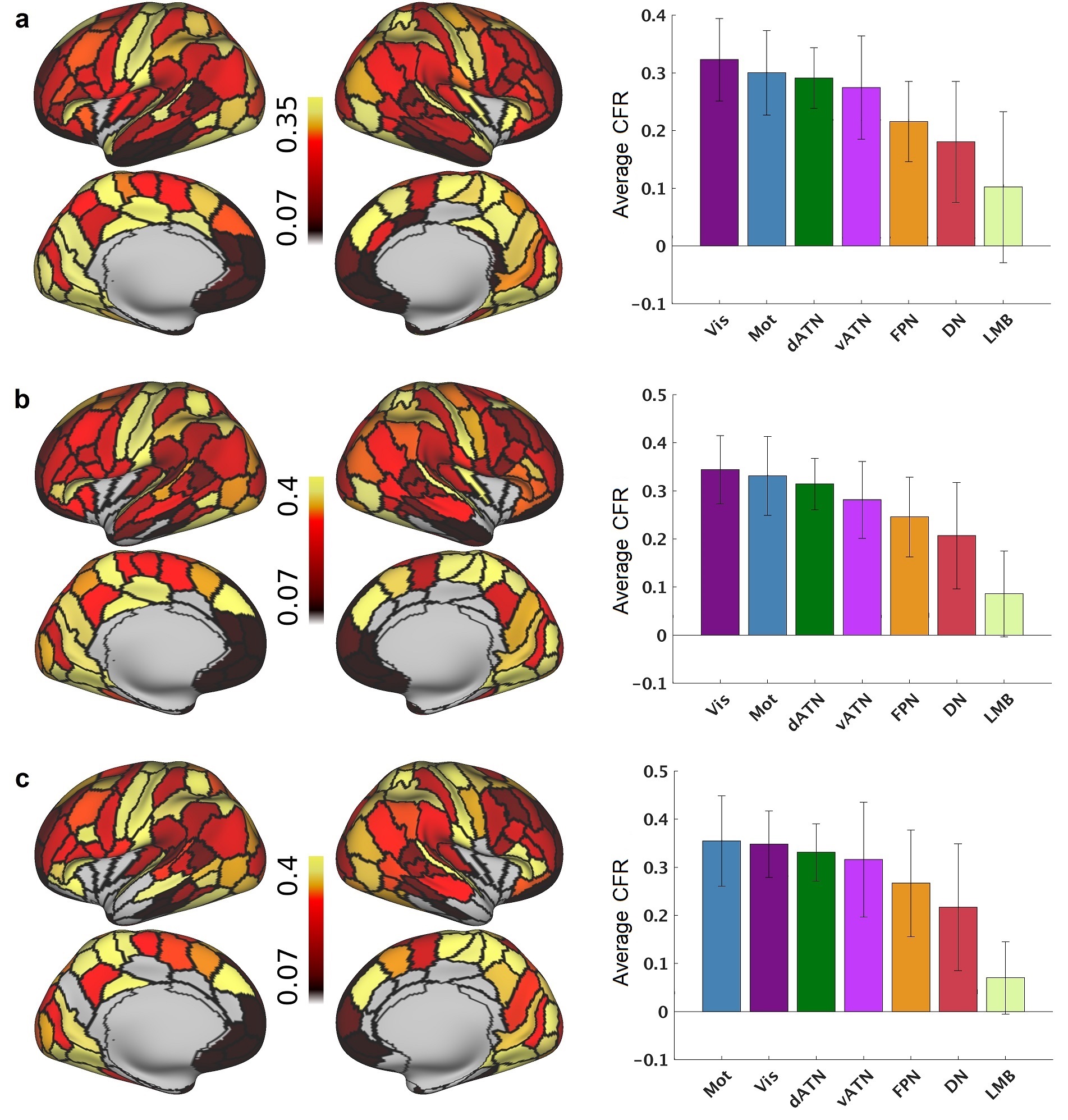

**Figure S3. Spatial distribution of the average CFR at different threshold levels.** Different threshold levels were used in the selective averaging process. The right endpoint of the 10%, 15% and 20% interval of the density peak was used as the threshold in (a), (b) and (c) respectively. The pattern of the spatial distribution of the average CFR was consistent under different threshold levels. The average CFR in the sensory-motor cortex is significantly (*p*<1e-6) higher than that in the association cortex under a two sample *t*-test.

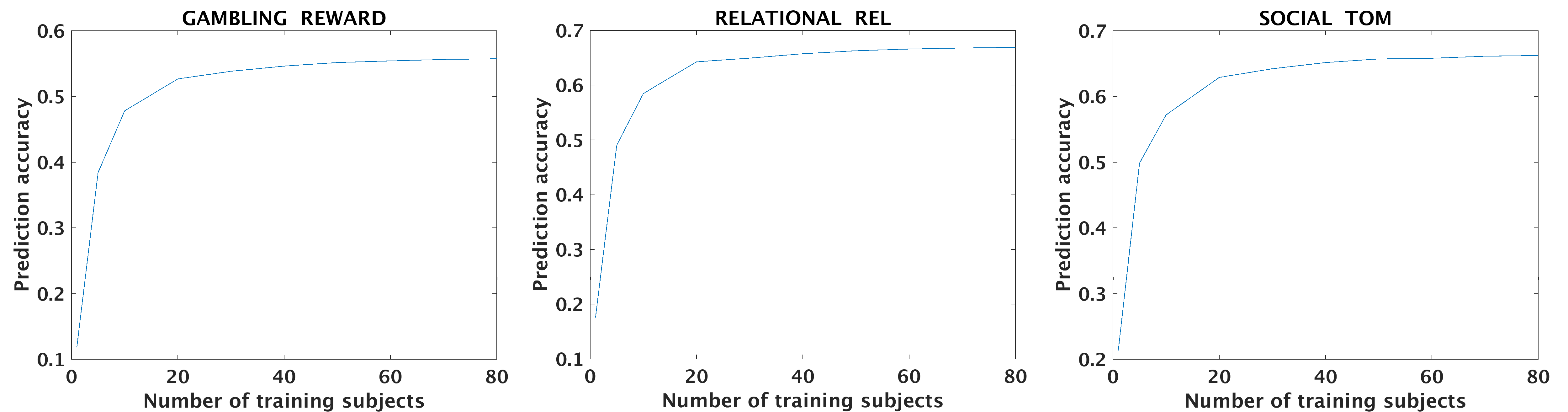

**Figure S4. The effect of training sample size on the prediction accuracy.** We tested the effect of training sample size on the prediction accuracy in the whole cortex and showed the results of three contrasts (other contrasts have similar results). We increased the number of training subject from 1 to 80 with the number of testing subject being 20. The model got only a little gain (less than 0.03) from increasing the sample size after 40 training subjects and the curves were nearly flat at 80 training subjects. Therefore, 80 training subjects and 20 testing subjects were enough to test the CFR accurately.

Subjects ID:

'100307'	'100408'	'101107'	'101309'	'101915'	'103111'	'103414'

'103818'	'105014'	'105115'	'106016'	'108828'	'110411'	'111312'

'111716'	'113619'	'113922'	'114419'	'115320'	'116524'	'117122'

'118528'	'118730'	'118932'	'120111'	'122317'	'122620'	'123117'

'123925'	'124422'	'125525'	'126325'	'127630'	'127933'	'128127'

'128632'	'129028'	'130013'	'130316'	'131217'	'131722'	'133019'

'133928'	'135225'	'135932'	'136833'	'138534'	'139637'	'140925'

'144832'	'146432'	'147737'	'148335'	'148840'	'149337'	'149539'

'149741'	'151223'	'151526'	'151627'	'153025'	'154734'	'156637'

'159340'	'160123'	'161731'	'162733'	'163129'	'176542'	'178950'

'188347'	'189450'	'190031'	'192540'	'196750'	'198451'	'199655'

'201111'	'208226'	'211417'	'211720'	'212318'	'214423'	'221319'

'239944'	'245333'	'280739'	'298051'	'366446'	'397760'	'414229'

'499566'	'654754'	'672756'	'751348'	'756055'	'792564'	'856766'

'857263'	'899885'

**Table S1. Subjects identification details.** The 100 unrelated subjects were random chosen from the HCP S1200 release.

| **Domain** | **Contrast** | **Similarity (Prediction)** | **Similarity**  **(Parcellation)** |
| --- | --- | --- | --- |
| EMOTION | FACES  SHAPES  FACES-SHAPES | 0.55  0.52  0.44 | 0.88  0.83  0.78 |
| GAMBLING | PUNISH  REWARD  PUNISH-REWARD | 0.56  0.58  0.11 | 0.86  0.86  0.53 |
| LANGUAGE | MATH  STORY  STORY-MATH | 0.40  0.47  0.59 | 0.82  0.86  0.86 |
| MOTOR | CUE  LF  LH  RF  RH  T  AVG  CUE-AVG  LF-AVG  LH-AVG  RF-AVG  RH-AVG  T-AVG | 0.61  0.40  0.37  0.39  0.35  0.42  0.41  0.60  0.32  0.40  0.37  0.33  0.39 | 0.89  0.78  0.73  0.78  0.74  0.79  0.78  0.89  0.75  0.78  0.74  0.72  0.73 |
| RELATIONAL | MATCH  REL  REL-MATCH | 0.68  0.69  0.26 | 0.90  0.90  0.66 |
| SOCIAL | RANDOM  TOM  TOM-RANDOM | 0.68  0.68  0.37 | 0.89  0.89  0.77 |
| WM | 2BK_BODY  2BK_FACE  2BK_PLACE  2BK_TOOL  0BK_BODY  0BK_FACE  0BK_PLACE  0BK_TOOL  2BK  0BK  2BK-0BK  BODY  FACE  PLACE  TOOL  BODY-AVG  FACE-AVG  PLACE-AVG  TOOL-AVG | 0.55  0.55  0.58  0.55  0.50  0.46  0.59  0.58  0.63  0.62  0.39  0.58  0.56  0.63  0.62  0.32  0.41  0.40  0.27 | 0.86  0.84  0.84  0.85  0.85  0.79  0.84  0.87  0.86  0.86  0.83  0.87  0.82  0.85  0.87  0.71  0.82  0.75  0.71 |

**Table S2. A summary table of similarity analysis.** The first column represents the seven task domains. The second column represents the 47 contrasts. The third column represents the mean similarity (evaluated by Pearson correlation) between each subject’s predicted activation and actual activation. The fourth column represents the mean similarity between the two predicted activations based on two different parcellations (Brainnetome and Multi-modal parcellation). The predictions based on the two parcellation were very similar and had very high correlation coefficients. Therefore, which kind of parcellation to use did not have a strong effect on the results.

| **Domain** | **Contrast** | **AC** | **AC-Distance** |
| --- | --- | --- | --- |
| EMOTION | FACES  SHAPES  FACES-SHAPES | 152  157  157 | 165  164  162 |
| GAMBLING | PUNISH  REWARD  PUNISH-REWARD | 178  182  49 | 181  185  56 |
| LANGUAGE | MATH  STORY  STORY-MATH | 163  149  194 | 177  162  194 |
| MOTOR | CUE  LF  LH  RF  RH  T  AVG  CUE-AVG  LF-AVG  LH-AVG  RF-AVG  RH-AVG  T-AVG | 174  133  117  135  123  126  147  179  85  90  76  85  94 | 176  143  129  140  131  135  158  176  88  93  92  80  104 |
| RELATIONAL | MATCH  REL  REL-MATCH | 185  188  92 | 188  187  103 |
| SOCIAL | RANDOM  TOM  TOM-RANDOM | 181  194  166 | 189  195  170 |
| WM | 2BK_BODY  2BK_FACE  2BK_PLACE  2BK_TOOL  0BK_BODY  0BK_FACE  0BK_PLACE  0BK_TOOL  2BK  0BK  2BK-0BK  BODY  FACE  PLACE  TOOL  BODY-AVG  FACE-AVG  PLACE-AVG  TOOL-AVG | 178  184  175  179  150  151  169  171  190  181  149  174  180  188  189  99  103  119  77 | 183  185  178  181  167  164  183  175  189  190  142  186  187  188  191  108  114  129  92 |

**Table S3. Statistical testing of conenctivity model.** We calculated the number of regions that had connectivity model statistically better than random in the column of AC (anatomical connectivity). We also calculated the number of regions that had connectivity model statistically better than random after regressing out distance in the column of AC-Distance. Anatomical connectivity had predictions statistically better than random in many regions across most contrasts, even after regressing out distance from the connectivity profile. And anatomical connectivity could even had predictions statistically better than random in a few more regions after regressing out euclidean distance. We did two sample *t*-tests and found that the regions that had predictions better than random had mean absolute activations significantly higher (*p*<0.001) than regions that had predictions no better than random, which indicated that it was most likely to have reliable predictions in high-activation regions. A few contrasts’ predictions such as the PUNISH-REWARD were better than random in only a few regions, because these contrasts had very low activation levels throughout the whole cortex.

| **Domain** | **Contrast** | **Task activation**(+) | | **Inter-subject task variation**(-) | |
| --- | --- | --- | --- | --- | --- |
|  |  | **CFR** | **CFR-variation** | **CFR** | **CFR-activation** |
| EMOTION | FACES  SHAPES  FACES-SHAPES | 0.16  0.11  0.43 | 0.05  0.09  0 | 0.60  0.46  0.67 | 0.29  0.19  0.13 |
| GAMBLING | PUNISH  REWARD  PUNISH-REWARD | 0.14  0.13  0.01 | 0.04  0.04  0 | 0.57  0.59  0.45 | 0.29  0.33  0.44 |
| LANGUAGE | MATH  STORY  STORY-MATH | 0.02  0.19  0.06 | 0.01  0.01  0.10 | 0.44  0.67  0.57 | 0.39  0.38  0.40 |
| MOTOR | CUE  LF  LH  RF  RH  T  AVG  CUE-AVG  LF-AVG  LH-AVG  RF-AVG  RH-AVG  T-AVG | 0.02  0.39  0.41  0.38  0.46  0.32  0.21  0.02  0.59  0.49  0.56  0.51  0.39 | 0.06  0.03  0.02  0.01  0.02  0.05  0.02  0.02  0.02  0.02  0.02  0.03  0.04 | 0.37  0.69  0.68  0.63  0.72  0.67  0.53  0.38  0.81  0.70  0.81  0.76  0.72 | 0.30  0.13  0.12  0.11  0.11  0.17  0.19  0.32  0.09  0.07  0.12  0.10  0.16 |
| RELATIONAL | MATCH  REL  REL-MATCH | 0.02  0.04  0.26 | 0.09  0.07  0.02 | 0.51  0.58  0.44 | 0.44  0.46  0.05 |
| SOCIAL | RANDOM  TOM  TOM-RANDOM | 0.10  0.05  0.19 | 0.02  0.03  0.07 | 0.55  0.53  0.64 | 0.36  0.40  0.28 |
| WM | 2BK_BODY  2BK_FACE  2BK_PLACE  2BK_TOOL  0BK_BODY  0BK_FACE  0BK_PLACE  0BK_TOOL  2BK  0BK  2BK-0BK  BODY  FACE  PLACE  TOOL  BODY-AVG  FACE-AVG  PLACE-AVG  TOOL-AVG | 0.13  0.19  0.33  0.14  0.22  0.32  0.34  0.33  0.07  0.28  0.07  0.11  0.24  0.29  0.26  0.51  0.32  0.52  0.44 | 0.02  0.04  0  0.06  0.02  0.04  0.01  0  0.06  0.01  0.02  0.01  0.02  0  0  0.01  0.07  0.02  0 | 0.51  0.61  0.61  0.63  0.63  0.74  0.67  0.66  0.56  0.64  0.48  0.44  0.65  0.63  0.61  0.81  0.78  0.80  0.78 | 0.27  0.26  0.18  0.35  0.27  0.25  0.20  0.22  0.39  0.24  0.32  0.23  0.27  0.24  0.25  0.19  0.29  0.15  0.30 |

**Table S4. Quantitative analysis of the factors that affeced the CFR based on Brainnetome atlas.** We calculated the amount of variance in the CFR that could be explained by task activation or inter-subject task variation. The plus sign indicated positive correlation with the CFR and the minus sign indicated negative correlation with the CFR. Task activation was positively correlated with the CFR, whereas inter-subject task variation was negatively correlated with the CFR. Task variation explained significantly (*p* < 1e-6) more variance in the CFR than task activation did under paired-*t* test across all contrasts. Task activation could barely explain no variance in the CFR after inter-subject task variation was regressed out, as indicated by the column of CFR-variation.
